# Whole-cortex simulation reveals spatiotemporal patterns emerging from the interplay of network connectivity and intracellular dynamics

**DOI:** 10.1101/2024.01.10.574958

**Authors:** Guanhua Sun, James Hazelden, Ruby Kim, Daniel Forger

## Abstract

Recent advances in Graphics Processing Unit (GPU) computing have allowed for computational models of whole-brain activity at unprecedented scales. In this work, we use desktop computers to build and simulate a whole-cortex mouse brain model using Hodgkin-Huxley type models for all the most active neurons in the mouse cortex. We compare the model dynamics over different types of connectivity, ranging from uniform random to realistic connectivity derived from experimental data on cell positions and the Allen Brain Atlas. By changing the external drive and coupling strength of neurons in the network, we can produce a wide range of oscillations in the gamma through delta bands. While the global mean-field behaviors of different connectivities share some similarities, an experimentally determined hierarchical connectivity allows for complex, heterogeneous behaviors typically seen in EEG recordings that are not observed in networks with nearest neighbors or uniform coupling. Moreover, our simulations reveal a wide range of spatiotemporal patterns, such as rotational or planar traveling waves, that are observed in experiments. Different traveling waves are observed with different connectivity and coupling strengths on the same connectivity. Our simulations show that many cortical behaviors emerge at scale with the full complexity of the network structure and ionic dynamics. We also provide a computational framework to explore these cortex-wide behaviors further.

## 1 Introduction

Researchers can measure the brain at scale, for example, with electroencephalography (EEG) or functional magnetic resonance imaging (fMRI) techniques[1–3]. In addition to EEG and fMRI, tremendous advances in single-cell spatial biology provide information on the physiology of individual cells within tissues[4, 5]. There is also a growing appreciation of the importance of spatiotemporal patterns in the brain, from traveling slow waves[6], which are the signature of deep sleep, to waves related to working memory[7], spiral waves during sleep spindles[8], and hemodynamic and cerebrospinal fluid waves that are coupled with electrophysiology[9]. These spatiotemporal patterns are crucial for understanding sleep, working memory, and other brain functions[10]. Yet how they arise from the collective behavior of single cells remains to be determined.

While simplified models such as firing-rate models[11] provide a way to understand some aspects of neuronal dynamics, direct simulation can help illuminate how the electrophysiology of individual cells contributes to complex network-level behaviors such as traveling waves. Realistic brain networks are known to be hierarchical[12, 13], but only in recent years have researchers conducted large-scale simulations with supercomputers to account for the complex structure[14, 15]. Although Hodgkin and Huxley[16] provided a detailed understanding of the electrical activity of a single neuron, the complexity of large-scale neuronal networks has made it difficult to extend this knowledge to whole-brain activity. Recent advances in computational power, mainly using computing on graphics processing units (GPUs), have made large-scale neuronal simulations on desktop computers plausible. Platforms such as GeNN[17], Neuron[18], Brian[19], TVB[20] and various studies[21, 22] have utilized GPU computing to simulate large-scale neuronal networks. However, these platforms still lack basic interfaces to construct or simulate complex connectivity, and there is also a need for real-time interactive visualization to observe possible spatiotemporal patterns happening during the simulations.

In a companion manuscript, we experimentally identified hundreds of thousands of active neurons within the mouse cortex. We mapped out their connections based on realistic connectivity data from the Allen Brain Atlas[23, 24]. This connectivity captures the cortex’s hierarchical structure(**Fig S1b**) but is still sparse because of the limited number of synapses on a neuron, making it possible to be simulated with a new framework.

In this work, we first develop a GPU-powered framework to simulate large-scale neuronal networks with interactive real-time visualization on a desktop GPU. We then use this framework to build different connectivities, such as uniform-random or local networks. Together with the realistic connectivity derived from the Allen Brain Atlas, we conduct extensive simulations of models with different connectivities. By varying the applied current to the neurons and synaptic strength between them, we can produce a range of oscillations, ranging from gamma to delta bands. While the global mean-field behaviors are similar for uniform-random or local connectivity, realistic connectivity creates more heterogeneity among different regions, resembling experimental measurements.

Moreover, in our simulations, the physiologically based connectivity can generate a rotational wave across the whole cortex, which has been experimentally observed. Local connectivity is also able to display planar waves that are observed through different experiments[25–27]. Notably, both kind of waves reproduced in our simulations match the experimental findings in [28]. We also study how coupling strength will affect those traveling waves qualitatively. These findings demonstrate how cortex-wide spatiotemporal patterns can emerge from the interplay between the intracellular dynamics and the network structure, and our computational framework can become an essential tool for visualizing and understanding brain activity at scale.

## 2 Results

### 2.1 Whole-cortex computational model

We first present a computational model of the mouse cortex with more than 400,000 active neurons at the beginning of the daylight. The model consists of three parts: electrophysiology, anatomy, and connectivity. To describe the electrophysiology of neurons, we choose a single-compartment conductance model of cortical neurons, whose behavior has been extensively studied in [29] (**Methods 4.1)**. We consider excitatory neurons with glutamatergic synapses (AMPA) and inhibitory GABAergic neurons. For anatomic information, neuron positional and regional information is experimentally measured in our companion manuscript, and they are aligned with the Allen Brain Atlas Common Coordinate Framework(ABA-CCF) [4]. The cell atlas provides the excitatory/inhibitory information of neurons[30]. Lastly, we construct four kinds of connectivity over the neurons:

1. **Allen connectivity (A)** is built with the realistic connectivity data provided by the Allen Brain Atlas and our experiments in a companion manuscript. Neurons are connected with different probabilities based on their regional information at the resolution of 100*μm*.
2. **Local connectivity (L)** is based on the distance between neurons. This manuscript adopts a nearest-neighbor coupling in which neurons communicate within a near region and uses the experimentally determined positions of all neurons.
3. **Uniform random connectivity (U)** is built by connecting neurons across the whole cortex with a uniform probability.
4. **Local+Allen connectivity (AL)** is the combination of Allen and local connectivity.

By adding the local connectivity to the Allen connectivity, we aim to fill in the local details that the Allen Brain Atlas does not provide due to the resolution of the experiments.

We have four different connectivities, resulting in four models we simulate and compare. The details of construction, storage, and simulation of the connectivity are described in **Methods 4.2**.

To make these simulations possible on realistic timeframes, we construct a computational platform utilizing GPUs for simulating and visualizing neural dynamics in real time. This framework encodes simulations by interacting modules, e.g., connectivity, spatial morphology of the network, and electrophysiology, to make the programming more accessible and readable. This framework allows us to use experimentally validated and detailed models of both intracellular electrophysiology and experimentally determined network structure. We explore how the interplay between network connectivity and nonlinear dynamics of individual neurons can induce a range of oscillations and spatiotemporal behaviors.

### 2.2 Changing neurons’ excitability and coupling strength can induce a range of global oscillations

In simulations, we study the effects of two main parameters: the applied current (I), which regulates the neuron’s excitability, and the coupling strength (g), which controls the magnitude of the synaptic current between neurons. For each type of connectivity, we vary the applied current from 0.6*μA* to 2.0*μA* and the conductance for AMPA receptors from 0.2 to 3.0*mS/cm*^2^ and study how they affect the network’s firing behavior and spatiotemporal dynamics. Looking at the mean-field behaviors of different models, we first find that all connectivities can generate a wide range of oscillations in gamma (30+ Hz), beta (13-30 Hz), alpha (8-12 Hz), theta (4-12 Hz), and even delta (0.5-4 Hz) bands (**Fig 2a**). These frequencies of neuronal firing correspond to different cortical functions such as motor tasks, learning, working memory, or sleep [31] [32]. For much of the parameter space, the dominant frequency of the average voltage across the whole cortex generally decreases from gamma to delta frequency bands as the coupling strength increases, indicating a transition to a sleep state (**Fig 2b**). In particular, at the same coupling strength, we see the gamma oscillations with the Allen connectivity have an outstanding amplitude compared to the other three connectivities, indicating a higher level of neuronal synchrony (**Fig 2b, Gamma row**).

**Fig 1:**
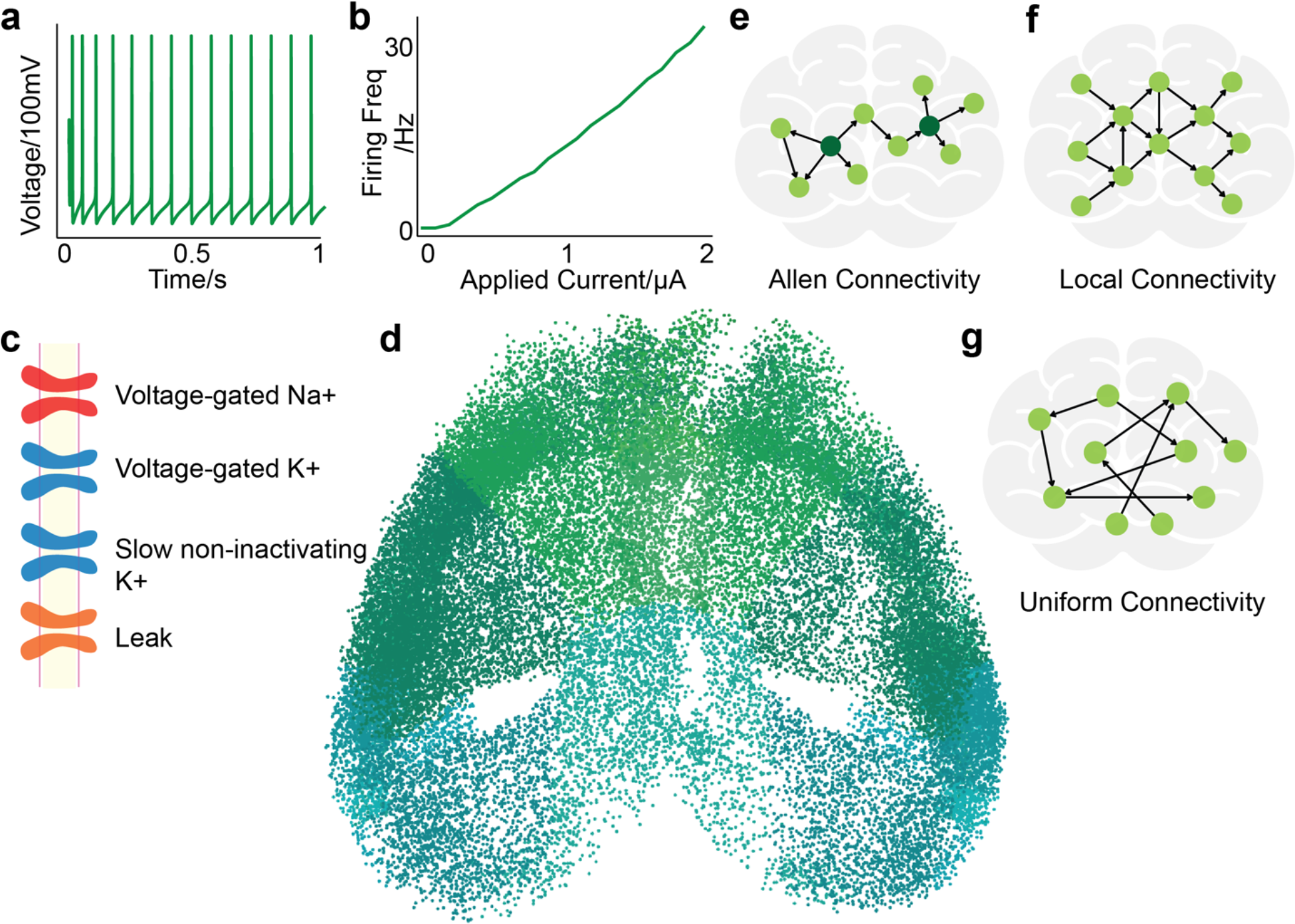
Overview of our computational model. **a-c:** The electrophysiology model of each neuron. **a:** Each neuron will fire regularly under applied current and does not intrinsically burst. **b:** The F-I curve of the neuron model. **c:** Demonstration of the membrane electrophysiology, illustrating the five ionic channels used to model each individual neuron. **d:** Horizontal view of the cortical neurons in our simulations. Each neuron is colored based on its region according to the Allen Brain Atlas[] colormap. **e-g:** Illustration of the three types of connectivities that we use in our simulation: Allen Brain connectivity that is based on the measured voxelized connectivity, local connectivity that is constructed based on the distance between neurons and uniform connectivity that span randomly across the whole-cortex.

**Fig 2:**
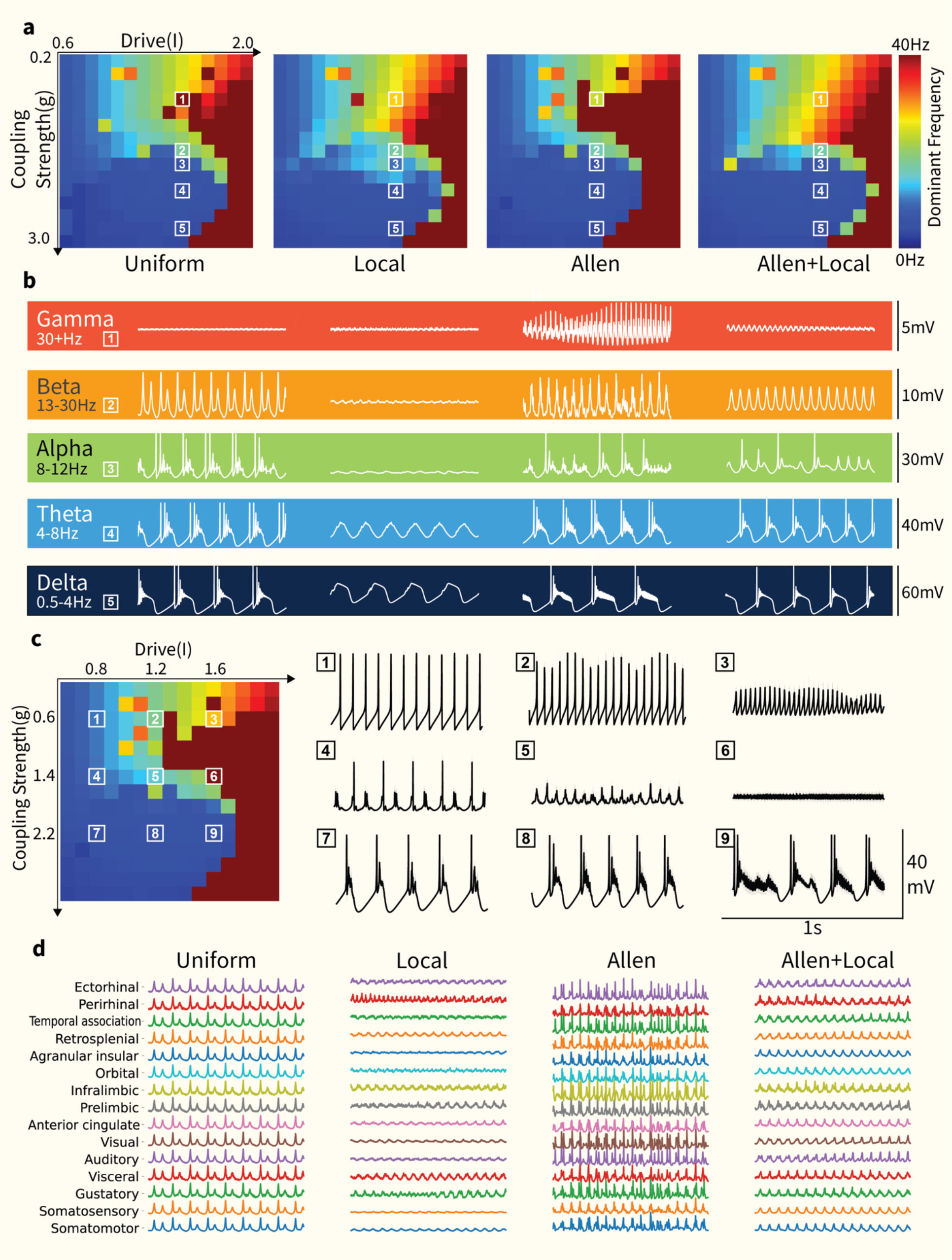
Variations of drive and synaptic coupling are able to produce different brain rhythms under different networks. **a**: Heat maps illustrating the dominant frequency of the mean voltage across the whole cortex under different levels of drive and excitatory coupling strength. Dominant frequency is measured as the frequency with strongest spectral power after spectrum analysis. Similar patterns are observed here for the different connectivities, indicating all connectivities result in similar global mean activity and are able to induce a range of different oscillations under different conditions. White boxes indicate the parameters corresponding to the recordings from (b). **b**: Plots of global mean voltage across the cortex with the different networks in (a) for 1s. Network bursting emerges in the presence of sufficiently strong excitatory coupling. The dominant frequency decreases when increasing the coupling strength. **Top to Bottom**: The excitatory synaptic coupling is increased (g=0.8, g=1.6, g=1.8, g=2.2, g=2.8 mS/cm^2), as indicated from 1-5 in (a). **Left to Right:** Uniform random (U), Local (L), Allen (A) and Allen+Local (AL) connectivity. **c**: Further analysis of the Allen heat map activity. We choose 9 sets of parameters from the heatmap**(left)** and plot the corresponding mean voltages**(right)**. Increase in drive makes the firing rate rise, but increase in coupling strength changes the firing behavior qualitatively. **d**: Local field potential (LFP) traces of major cortical regions for 1s. This particular set of simulations corresponds to the parameters of box (2) indicated in figure (a). Uniform and local connectivity results in highly uniform LFP across different areas, whereas other connectivity that contain the Allen connectivity patterns produce more heterogeneity.

The frequency transitions can also be non-monotonic as coupling strength is increased. For all four connectivity models and some values of *I*, there is an increase in frequency before an abrupt transition to lower frequency ranges. Then, as coupling strength is increased even further, the dominant frequency “jumps” back to higher ranges (**Fig 2a**).

### 2.3 Interplay between recurrent excitation and inhibitory ionic currents causes network bursting

Both the whole-cortex average and LFP recordings in our simulations demonstrate a transition to bursting when the strength of the excitatory coupling increases. Neuronal populations that demonstrate bursting while individual neurons spike is known as *network bursting* and is a known behavior of neuronal networks with excitatory and inhibitory neurons [33] [34] [35]. As shown in **Fig 2c**, if we fix the coupling strength but only increase the applied current to each neuron, we see that we merely increase the firing rate of the system but do not change the qualitative behavior of the firing (**Fig 2c, 1-2-3**). On the other hand, if we fix the applied current and increase the coupling strength, the network transitions from regular firing to network bursting(**Fig 2c 1-4-7**). Furthermore, the number of firings per burst and the interval between each burst are related to the strength of the coupling and the strength of the drive.

Counterintuitively, the dominant frequency of the oscillation decreases with increasing coupling strength. As a result, delta waves happen at the strongest coupling strength. This may be important for slow waves in sleep research because the interplay between network bursting and silence resembles the up-down dynamics of slow-wave activity(SWA), the electrophysiology hallmark of NREM sleep [36]. It also has been shown that the strength of SWA depends on the coupling strength [37], which is reaffirmed by our simulations.

### 2.4 Realistic connectivity creates heterogeneous regional recordings

Though the phase plots in **Fig 2a** share some qualitative similarities across the four connectivities, they plot the dominant frequency of the average global electrical activity across the whole cortex. Thus, they do not tell the full story of the spatiotemporal dynamics. To dive deeper into the firing activity, we measure the local field potential at 15 major cortical subregions, such as visual, auditory, and somatosensory cortex. Realistic Allen connectivity produces more regional heterogeneity than uniform and local connectivity(**Fig 2d**) for beta and alpha oscillations. This finding demonstrates that the hierarchy and heterogeneity in the Allen connectivity can create regionally heterogeneous firing behaviors that closely resemble the experiments. Surprisingly, the Allen+Local connectivity did not show as much regional heterogeneity compared to Allen connectivity, indicating that the local connectivity may dominate in this simulation.

### 2.5 Realistic connectivity induces rotational traveling waves

We next explore the spatio-temporal patterns at different oscillation frequencies we observe with different drive and coupling strengths with the Allen connectivity. From our visualizations of the simulations, we found that the firing population travels rotationally across the whole cortex (**Fig 3a, Movie S1**). The rotational structure is better captured by coloring neurons based on the time they fire during a period(**Fig 3b**). Such global rotational traveling waves have been observed in the cortex [28] during tasks such as working memory or sleep, particularly sleep spindles. Unlike most past computational models that managed to reproduce rotational dynamics using a network of Kuramoto oscillators [38] with local coupling [39], the Allen connectivity we use here is a long-range global and long-range connectivity. This is the first time such rotational waves have been observed with realistic physical connectivity. Our findings here show that the rotational dynamics can emerge from the hierarchy of the network and the nonlinear dynamics of each neuron, which indicates that the rotational dynamics may have a deeper connection with the cortical structure.

**Fig 3:**
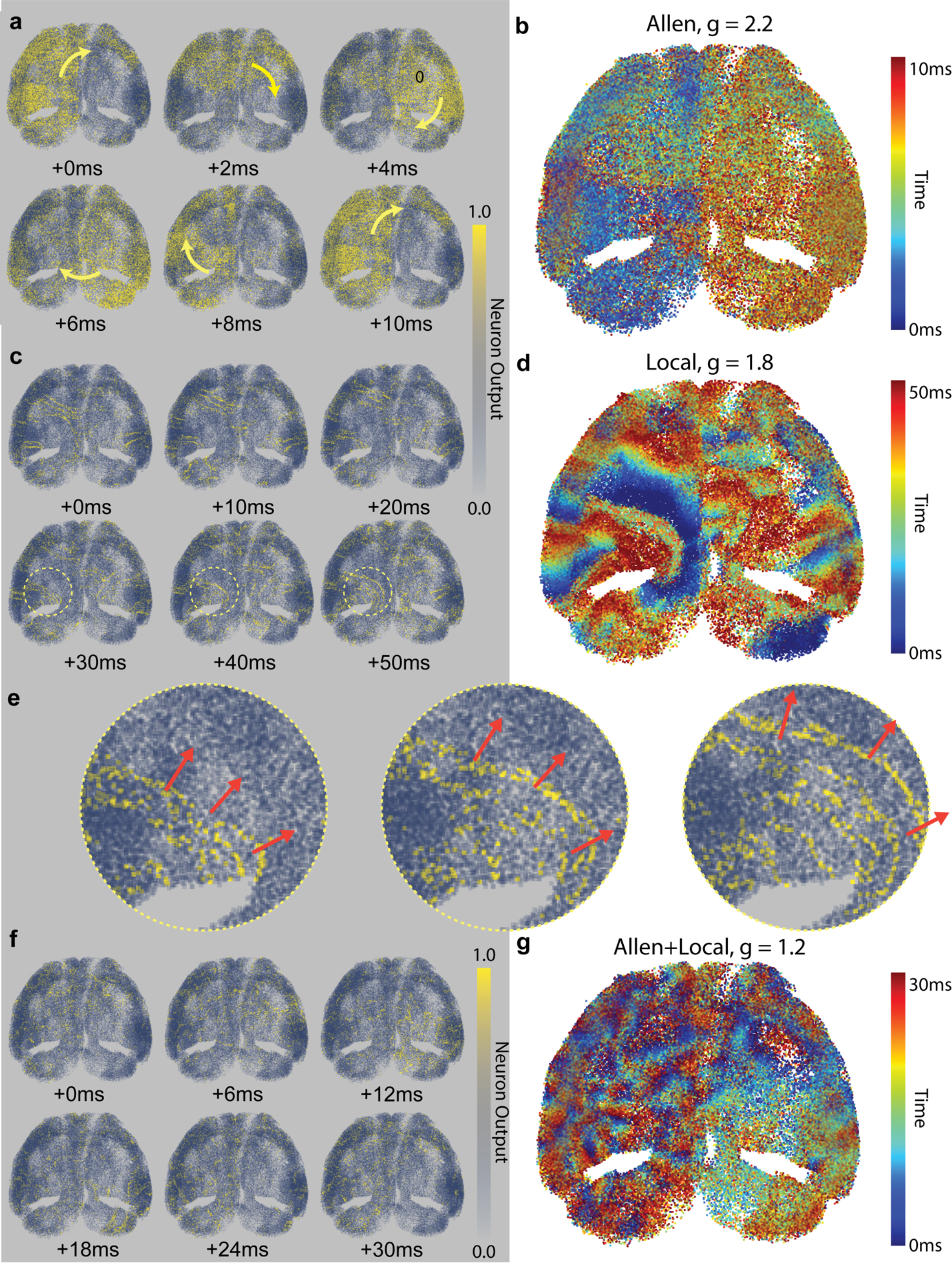
Different traveling waves induced by different network connectivity. **a-b:** Rotational waves induced by Allen connectivity. **a:** Each dot represents the synaptic output of a neuron at that position, which is a sigmoidal function of the voltage (see Methods 4.2). We see the firing pattern travels around the cortex rotationally. **b:** The activity in (a) is visualized by coloring neurons based on the time at which they fire during the span of 10 milliseconds. The overall wheel-shape pattern shows the rotational dynamics. **c-e:** Planar traveling waves induced by local connectivity (L). **c:** Action potentials extend radially in planar traveling waves. **d:** As in (b), neurons are colored based on the firing time in the time span of 50 milliseconds, illustrating action potential propagation as planar waves. **e:** Close-up of the dotted circled region in (d), where firing activities propagate during the time window of 20ms. **f-g**: Mixed traveling waves induced by Allen+local (AL) connectivity. **f:** Here, we see a mixing of the rotational gradation observed in (a) combined with the local traveling waves of (b). **g:** 30 millisecond temporal coloring, as in (b) and (d).

### 2.6 Local connectivity induces planar traveling waves

At the same time, local connectivity can induce another kind of traveling wave seen in our simulations: planar waves. Such waves are depicted in (**Figs 3c&e, Movie S2**), where the traveling wave usually starts from a small group of neurons firing and further expands to adjacent regions. Since planar traveling waves are not periodic, we visualize the propagation of those waves during 50ms in **Fig 3d**. Planar traveling waves are more frequently observed in different brain regions during other tasks and at different frequencies [26] [27]. Various computational models have obtained similar results, again utilizing Kuramoto oscillators with local connectivity[40]. However, in most of those studies, the structure of local connectivity is comparable to that of the global scale. In our simulations, the scale of local connectivity is far more localized, thus creating the opportunity for different waves to interact and produce more complicated recurrent patterns.

### 2.7 Coexistence of global and local dynamics

Next, we look at the results of Allen+Local connectivity to investigate how local and global dynamics can interact. In simulations with the combined connectivity, we observe that local dynamics can coexist with global dynamics. As shown in (**Fig 3f-g)**, we can combine global rotational traveling and local planar waves. A rotational envelope develops globally in those simulations, while local waves can also propagate inside the larger envelope(**Movie S3**). Again, the scale of our simulations is enough to contain the coexistence of global and local dynamics. As discussed in [41], such coexistence may only appear when the scale of the model is large enough.

Interestingly, when we measure the global synchrony and compare the case with Allen connectivity only and Allen+Local connectivity, we find that the latter case always has lower global synchrony. Such findings show that strong local connectivity may not help the whole network to synchronize(**Fig S2)**. The local planar waves may disturb the high level of synchrony that is achieved with global connections.

### 2.8 Traveling waves are affected by network structure and coupling strength

Lastly, we want to see how the coupling strength that controls the recurrent activity within the network affects different properties of traveling waves. By looking at the spatio- temporal patterns of different oscillations, we find that all oscillations under uniform-random connectivity do not appear to have any particular spatial pattern **Fig 4U(1-5)**: the firing activities are mostly synchronized across the cortex. Next, planar traveling waves are observed in all simulations with local connectivity, but the coupling strength affects the wavelengths and speeds. As shown in **Fig 4L(1-5),** the wavelength largely increases as the coupling strength increases while the dominant frequency decreases. The overall spatio-temporal dynamics structure with the Allen connectivity seems unchanged, as seen in **Fig 4A(1-5)**. The rotational dynamics of firing activities remain intact when changing the coupling strength, but the rotations’ period changes. Lastly, activities under Allen+Local change dramatically during theta and delta oscillations, compared to activities during gamma, beta, and alpha oscillations **Fig 4AL(1-5)**. Recall that the theta and delta oscillations are largely due to the network bursting. Such change shows that there could be a deeper connection between the oscillations and the network structure that dominates the dynamics.

**Fig 4:**
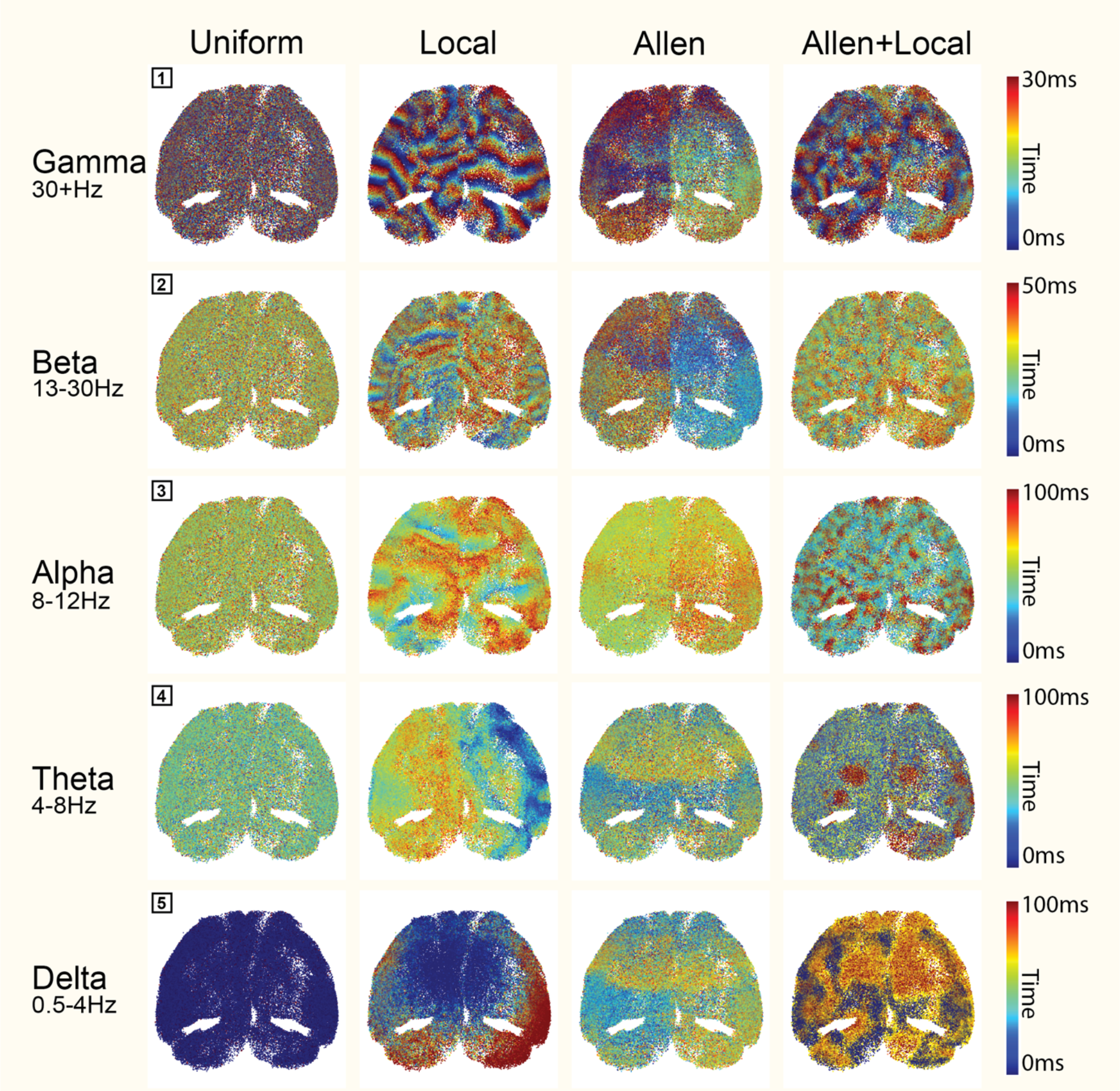
Spatiotemporal patterns of different oscillations with different networks. Temporal coding of firing activities over different time windows with four types of connectivity. Simulations marked with black boxes here **(1-5)** correspond to the simulations **(1-5)** for each connectivity presented in Fig 2b, where I = 1.4 *μA*. **Left to Right:** Uniform, local, Allen and Allen+Local connectivity. **Top to Bottom:** Increasing coupling strength(g=0.8, g=1.6, g=1.8, g=2.2, g=2.8 mS/cm^2) but decreasing dominant frequencies. This relationship is given by Fig2a.

## 3 Discussion

### 3.1 Recurrent activity and traveling waves

In this work, we develop a simulation framework to study the mouse cortex at scale. We find that the specific connectivity of the neuronal network can significantly influence the behaviors seen. An electrophysiology hallmark in our simulation is the network bursting behavior starting from alpha, and especially at the theta and delta bands. In those simulations where the network bursting is present, there are two states: a short period where neurons are firing and a silent period where all neurons are silent. The two states here resemble the up-down dynamics that constitute the slow-wave activity [42]. Notably, in our simulations, we only change the coupling strength of the excitatory synapses. Still, not the inhibitory ones, therefore changing the E/I balance of the network simultaneously, which also fits the explanation for the up-down dynamics. As excitatory synapses become stronger in the network, the increasing recurrent excitation can balance with the inhibition within the network, which is regulated by the slow-potassium current as an adaptation [42]. Potassium channels are important in generating and controlling slow waves [43] [44]. How the balance between recurrent excitation and inhibition affects the up-down dynamics has been extensively studied in [45] [46]. While these results match our simulations, we have revealed the spatio-temporal of those recurrent activities on different networks. In particular, the rotational and planar traveling waves may have further physiological implications not discussed in this paper.

### 3.2 Quantitative measurements of traveling waves

We show how realistic connectivity and intracellular electrophysiology can generate diverse traveling wave behaviors as connectivity, coupling strength, and external drive vary. Although our study primarily discusses behaviors qualitatively, extensive quantitative evaluations of these waves could yield more insights. Previous investigations have determined ways to characterize these waves, including measurement of wave speed and wavelength [26] [40]. In our experiments, the emergence of local planar waves, noted for their capacity to interact with adjacent waveforms, manifests complex, recurrent patterns. Accurately measuring parameters such as wave speed and length presents significant challenges in this context, necessitating the implementation of both artificial tagging of these waves and the development of sophisticated algorithmic measurement techniques. Despite these challenges, quantifying these wave properties remains a crucial objective for future research, offering potential alignment with experimental data and contributing to the broader understanding of wave dynamics in the cortex.

### 3.3 Understanding the cortex at scale with direct simulations

Different mathematical models have contributed significantly to understanding brain dynamics. Mean-field approximations, such as firing rate models, can accurately depict the average behavior of neuronal populations to an extent. As presented by this manuscript, although the individual neurons and local field potentials act very differently and the spatiotemporal patterns of different networks are different, the phase plots of the global average still share some similarities. Traveling waves have also been reproduced with mesoscopic models such as neural-mass or neural-field models [47]. However, direct simulations of large-scale neuronal networks will reveal unparalleled details and insights into brain dynamics in the future. The reason is twofold: first, we need more mathematical techniques to integrate the network structure into simplified models, such as firing rate models, built on mean-filed assumptions. Second, neurons’ highly nonlinear nature seems sensitive to network structure. Even in a much-simplified model of Kuramoto oscillators with uniform random connections, the global synchrony is very sensitive to the percentage of connections drawn through the network [48]. Moreover, different timescales of different ionic channels and synapses can interact with the network structure, as seen in this paper. Such phenomena will be challenging to reproduce with simple oscillators such as Kuramoto models or leak-integrate-fire neurons.

### 3.4 Simulation Platform

Our framework rapidly simulates hundreds of thousands of realistic point neurons with realistic network structures on a desktop GPU. However, this model is still far from a complete description of the cortex. Factors such as different regional electrophysiology, compartments of a single neuron, and synaptic delays are not accounted for. We also focus on the most active neurons experimentally determined in our companion manuscript at a particular time of day. Other times of the day or experimental conditions could yield a different connectivity with different neurons. Since we are interested in whole-cortex dynamics, we need to describe the local microcircuits in the cortex accurately. Measuring these throughout the cortex is a daunting task that may take decades to do. However, it could be incorporated into our framework if it were done. Neurons that are less active and not picked up by our cFos experiments may also play a role in behaviors. A novelty of our simulation framework is the efficient low-level visualization and the modular nature of the code, e.g., simulating EEG and fMRI. Additional modules could be included, providing a robust and interactive tool that computational neuroscientists could efficiently utilize.

## 4 Methods

### 4.1 Details of electrophysiology

We use a minimal Hodgkin-Huxley-type neuron model that can produce different firing behaviors analogous to a cortical neuron [29]. Each neuron has four ionic channels: voltage-gated sodium, voltage-gated potassium, slow non-inactivating potassium, and leak channels. We refer to the original paper for detailed equations of each current and the change of gating variables are provided in the supplements. The parameters of the neuronal model are also drawn from [29].

### 4.2 Connectivity and Coupling

Four types of connectives are used in our simulations: Allen, Local, Uniform, and Allen+Local connectivity. Please refer to the companion manuscript for descriptions of the Allen connectivity, which is based on the realistic connectivity data provided by Allen Brain Atlas. The local connectivity in our model is nearest-neighbor coupling, where we use the KNN algorithm [49] to decide a certain number of nearest neighbors for each neuron based on neurons’ positions. Uniform connectivity is constructed by selecting random downstream neurons through the cortex with equal probabilities. To note, both our local and uniform connectivity are fixed-degree networks, and the average number of downstream connections of each neuron is chosen to match the average produced by the Allen connectivity, which is 658 connections per neuron. Finally, Allen+Local connectivity combines Allen and local connectivity, where each neuron, on average, projects 2*658 = 1316 neurons.

Overall, each connectivity contains about 300 million synapses. Hence, parallelism is required to facilitate the simulation of such large networks. Neuron output is modeled as a sigmoidal function and all neurons are conductance-coupled. One approach to solving connectivity is propagating this signal to downstream neurons by sparse matrix-vector multiplication algorithm. However, similar to prior work [19] [50], we find that handling connectivity using an *adjacency list* representation is more efficient. Our approach determines which neurons are firing and then propagates synaptic signals by a weighted sum over these neurons’ outputs. These operations can be implemented in parallel and make connectivity much faster since the average number of neurons spiking at any instant is usually much lower than the total neuron count.

### 4.3 Numerical method and GPU parallelism

We solve the Hodgkin-Huxley equations using an implicit leapfrog rule [21], allowing for large timesteps due to increased stability. During simulation, electrophysiology and connectivity are simulated on the same time scale to avoid missing spikes, which is also used to judge whether a neuron is firing when processing synaptic connections. Both electrophysiology and synaptic equations are paralleled on GPU in our framework. The overall parallelism is based on the neuron, which means each set of Hodgkin-Huxley equations of a neuron is put up on an individual thread of GPU to solve at every timestep. Similarly, at any time step, we first check for all neurons firing and then assign each thread to process each of the firing neurons’ downstream connections.

### 4.4 Simulation and data collection

Each model presented in this paper is simulated for 30 seconds with our framework and all data are obtained with a sampling frequency of 2000Hz. Phase plots in **Fig 2a** are calculated based on the data of the last 5 seconds of the 30-second simulations. The global average voltage is calculated by calculating the average of 10,0000 randomly sampled neurons, about 25% of the total neurons. The local field potential of different regions is calculated by calculating the average voltage of neurons tested in those regions, following the regional information provided by the Allen Brain Atlas [4].

### 4.5 Visualization

We implement real-time, customizable dynamics visualization as part of our simulation framework. The primary means of visualization is through a colored 3D point cloud to represent the physical positions of neurons. Unlike other high-level graphic interfaces, we write our visualization module with low-level OpenGL language that directly interfaces with the simulation while running.

## Acknowledgments

We thank Ningyuan Wang for a helpful discussion about work and coding. This work is dedicated to Steven Brown, who provided many helpful suggestions.

We acknowledge the following funding: HFSP RGP0019/2018, NSF DMS 2052499, and ARO MURI W911NF-22-1-0223 to DBF.

---

**Fig S1:**
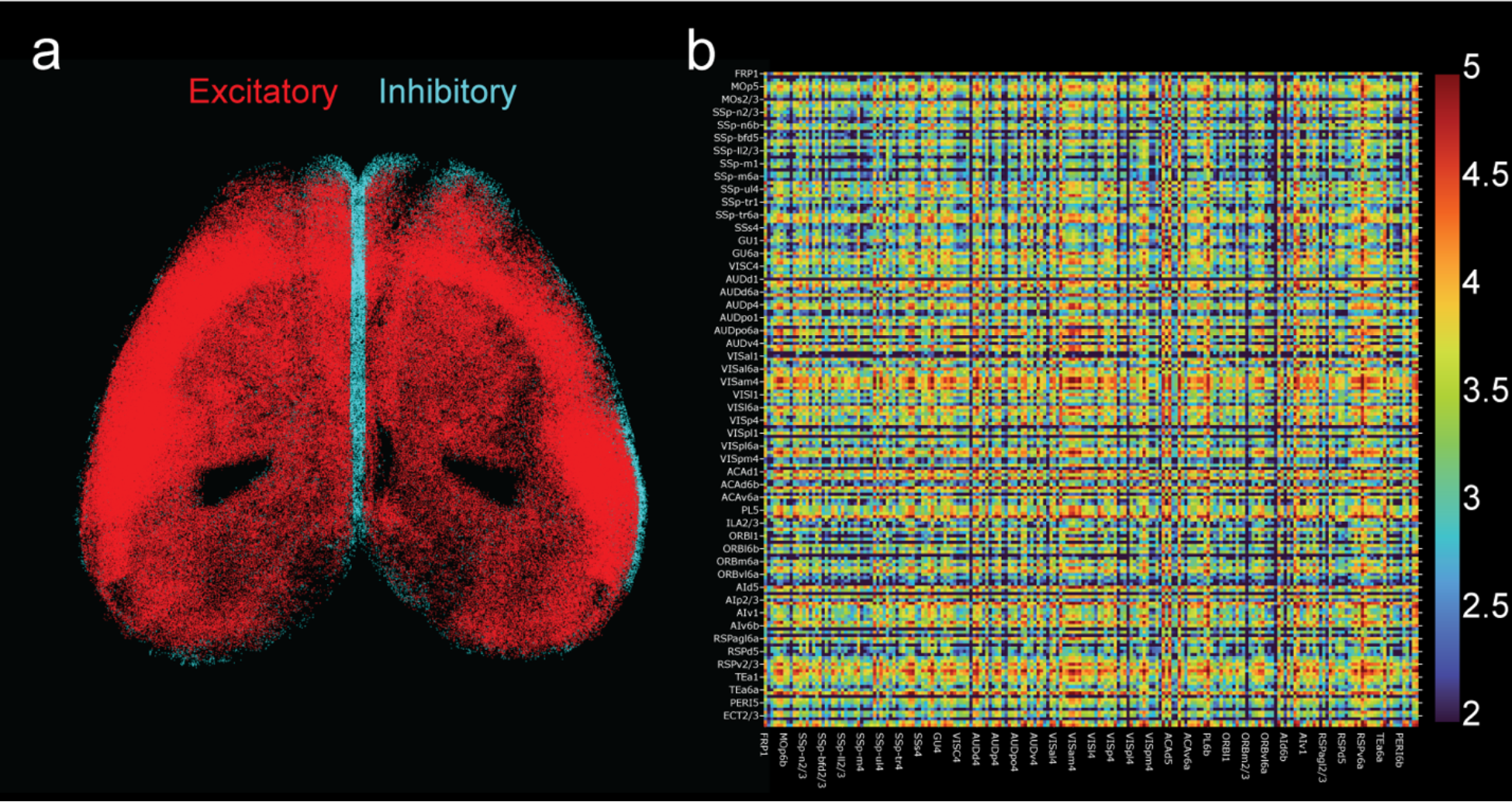
E/I information and regional connectivity matrix. **Left:** All cortical neurons visualized based on excitatory/inhibitory information. **Right**: The regional connectivity matrix. The cortical model we used has 205 subregions and the heatmap shows the number of connections between each region in logarithmic(log10) scale.

**Fig S2:**
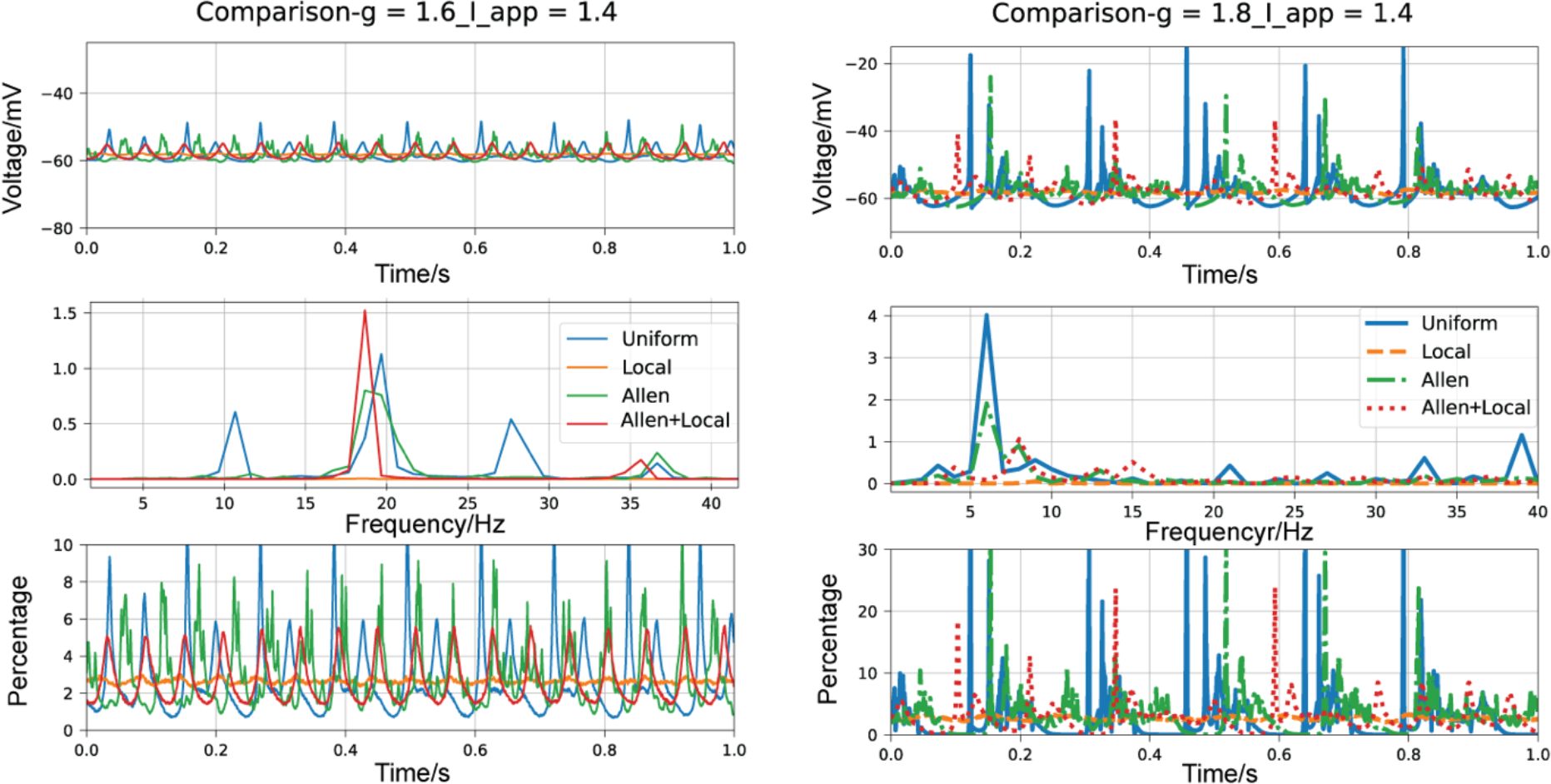
Comparison between the four connectivity for alpha and beta oscillations. Top: The average voltage of the whole cortex, where higher amplitude indicates a higher synchrony of neurons firing. **Mid:** The power spectrum of the average voltage presented above. **Bottom:** The percentage of neurons that are firing over the total population, where higher amplitude indicates a higher synchrony. Note all Allen+Local connectivity has lower synchrony than Allen connectivity alone.

## References

1. Michel CM, Koenig T. EEG microstates as a tool for studying the temporal dynamics of whole-brain neuronal networks: A review. Neuroimage. 2018;180(Pt B):577–93.

2. Markicevic M, Savvateev I, Grimm C, Zerbi V. Emerging imaging methods to study whole-brain function in rodent models. Translational Psychiatry. 2021;11(1):457.

3. Power JD, Plitt M, Laumann TO, Martin A. Sources and implications of whole-brain fMRI signals in humans. Neuroimage. 2017;146:609–25.

4. Wang Q, Ding S-L, Li Y, Royall J, Feng D, Lesnar P, et al. The Allen Mouse Brain Common Coordinate Framework: A 3D Reference Atlas. Cell. 2020;181(4):936–53.e20.

5. Murakami TC, Mano T, Saikawa S, Horiguchi SA, Shigeta D, Baba K, et al. A three-dimensional single-cell-resolution whole-brain atlas using CUBIC-X expansion microscopy and tissue clearing. Nature Neuroscience. 2018;21(4):625-+.

6. Massimini M. The Sleep Slow Oscillation as a Traveling Wave. Journal of Neuroscience. 2004;24(31):6862–70.

7. Bhattacharya S, Brincat SL, Lundqvist M, Miller EK. Traveling waves in the prefrontal cortex during working memory. PLoS Comput Biol. 2022;18(1):e1009827.

8. Muller L, Piantoni G, Koller D, Cash SS, Halgren E, Sejnowski TJ. Rotating waves during human sleep spindles organize global patterns of activity that repeat precisely through the night. Elife. 2016;5.

9. Fultz NE, Bonmassar G, Setsompop K, Stickgold RA, Rosen BR, Polimeni JR, et al. Coupled electrophysiological, hemodynamic, and cerebrospinal fluid oscillations in human sleep. Science. 2019;366(6465):628-31.

10. Muller L, Chavane F, Reynolds J, Sejnowski TJ. Cortical travelling waves: mechanisms and computational principles. Nat Rev Neurosci. 2018;19(5):255–68.

11. Wilson HR, Cowan JD. Excitatory and inhibitory interactions in localized populations of model neurons. Biophys J. 1972;12(1):1–24.

12. Harris JA, Mihalas S, Hirokawa KE, Whitesell JD, Choi H, Bernard A, et al. Hierarchical organization of cortical and thalamic connectivity. Nature. 2019;575(7781):195-+.

13. Coletta L, Pagani M, Whitesell JD, Harris JA, Bernhardt B, Gozzi A. Network structure of the mouse brain connectome with voxel resolution. Sci Adv. 2020;6(51).

14. Yamaura H, Igarashi J, Yamazaki T. Simulation of a human-scale cerebellar network model on the k computer. Front Neuroinform. 2020;14:16.

15. Markram H, Muller E, Ramaswamy S, Reimann Michael W, Abdellah M, Sanchez Carlos A, et al. Reconstruction and Simulation of Neocortical Microcircuitry. Cell. 2015;163(2):456–92.

16. Hodgkin AL, Huxley AF. A quantitative description of membrane current and its application to conduction and excitation in nerve. J Physiol. 1952;117(4):500–44.

17. Knight JC, Komissarov A, Nowotny T. PyGeNN: A Python Library for GPU-Enhanced Neural Networks. Front Neuroinform. 2021;15.

18. Hines ML, Carnevale NT. NEURON: a tool for neuroscientists. Neuroscientist. 2001;7(2):123–35.

19. Stimberg M, Brette R, Goodman DFM. Brian 2, an intuitive and efficient neural simulator. eLife. 2019;8:e47314.

20. Sanz Leon P, Knock SA, Woodman MM, Domide L, Mersmann J, McIntosh AR, et al. The Virtual Brain: a simulator of primate brain network dynamics. Front Neuroinform. 2013;7:10.

21. DeWoskin D, Myung J, Belle MDC, Piggins HD, Takumi T, Forger DB. Distinct roles for GABA across multiple timescales in mammalian circadian timekeeping. P Natl Acad Sci USA. 2015;112(29):E3911–E9.

22. Myung J, Hong S, DeWoskin D, De Schutter E, Forger DB, Takumi T. GABA-mediated repulsive coupling between circadian clock neurons in the SCN encodes seasonal time. Proceedings of the National Academy of Sciences. 2015;112(29):E3920–E9.

23. Oh SW, Harris JA, Ng L, Winslow B, Cain N, Mihalas S, et al. A mesoscale connectome of the mouse brain. Nature. 2014;508(7495):207-14.

24. Knox JE, Harris KD, Graddis N, Whitesell JD, Zeng HK, Harris JA, et al. High-resolution data-driven model of the mouse connectome. Netw Neurosci. 2018;3(1):217–36.

25. Lubenov EV, Siapas AG. Hippocampal theta oscillations are travelling waves. Nature. 2009;459(7246):534-9.

26. Zhang H, Jacobs J. Traveling Theta Waves in the Human Hippocampus. The Journal of Neuroscience. 2015;35(36):12477–87.

27. Sato TK, Nauhaus I, Carandini M. Traveling waves in visual cortex. Neuron. 2012;75(2):218–29.

28. Huang X, Xu W, Liang J, Takagaki K, Gao X, Wu JY. Spiral wave dynamics in neocortex. Neuron. 2010;68(5):978–90.

29. Pospischil M, Toledo-Rodriguez M, Monier C, Piwkowska Z, Bal T, Frégnac Y, et al. Minimal Hodgkin–Huxley type models for different classes of cortical and thalamic neurons. Biological Cybernetics. 2008;99(4-5):427–41.

30. Erö C, Gewaltig M-O, Keller D, Markram H. A Cell Atlas for the Mouse Brain. Front Neuroinform. 2018;12.

31. Wang XJ. Neurophysiological and computational principles of cortical rhythms in cognition. Physiol Rev. 2010;90(3):1195–268.

32. Buzsáki G, Draguhn A. Neuronal oscillations in cortical networks. Science. 2004;304(5679):1926-9.

33. Fardet T, Ballandras M, Bottani S, Métens S, Monceau P. Understanding the Generation of Network Bursts by Adaptive Oscillatory Neurons. Front Neurosci. 2018;12:41.

34. Masquelier T, Deco G. Network Bursting Dynamics in Excitatory Cortical Neuron Cultures Results from the Combination of Different Adaptive Mechanism. PLOS ONE. 2013;8(10):e75824.

35. Bogaard A, Parent J, Zochowski M, Booth V. Interaction of Cellular and Network Mechanisms in Spatiotemporal Pattern Formation in Neuronal Networks. Journal of Neuroscience. 2009;29(6):1677–87.

36. Amzica F, Steriade M. Electrophysiological correlates of sleep delta waves. Electroencephalography and clinical neurophysiology. 1998;107(2):69–83.

37. Esser SK, Hill SL, Tononi G. Sleep homeostasis and cortical synchronization: I. Modeling the effects of synaptic strength on sleep slow waves. Sleep. 2007;30(12):1617–30.

38. Kuramoto Y, Kuramoto Y. Chemical turbulence: Springer; 1984.

39. Ermentrout GB, Kleinfeld D. Traveling Electrical Waves in Cortex: Insights from Phase Dynamics and Speculation on a Computational Role. Neuron. 2001;29(1):33–44.

40. Zhang H, Watrous AJ, Patel A, Jacobs J. Theta and alpha oscillations are traveling waves in the human neocortex. Neuron. 2018;98(6):1269–81. e4.

41. Budzinski RC, Nguyen TT, Doan J, Minac J, Sejnowski TJ, Muller LE. Geometry unites synchrony, chimeras, and waves in nonlinear oscillator networks. Chaos. 2022;32(3):031104.

42. Compte A, Sanchez-Vives MV, McCormick DA, Wang XJ. Cellular and network mechanisms of slow oscillatory activity (< 1 Hz) and wave propagations in a cortical network model. Journal of Neurophysiology. 2003;89(5):2707–25.

43. Yoshida K, Shi S, Ukai-Tadenuma M, Fujishima H, Ohno RI, Ueda HR. Leak potassium channels regulate sleep duration. Proc Natl Acad Sci U S A. 2018;115(40):E9459–E68.

44. Muheim CM, Spinnler A, Sartorius T, Durr R, Huber R, Kabagema C, et al. Dynamic- and Frequency-Specific Regulation of Sleep Oscillations by Cortical Potassium Channels. Current Biology. 2019;29(18):2983-+.

45. Levenstein D, Buzsáki G, Rinzel J. NREM sleep in the rodent neocortex and hippocampus reflects excitable dynamics. Nature Communications. 2019;10(1).

46. Jercog D, Roxin A, Bartho P, Luczak A, Compte A, de la Rocha J. UP-DOWN cortical dynamics reflect state transitions in a bistable network. Elife. 2017;6.

47. Cakan C, Dimulescu C, Khakimova L, Obst D, Flöel A, Obermayer K. Spatiotemporal patterns of adaptation-induced slow oscillations in a whole-brain model of slow-wave sleep. Frontiers in computational neuroscience. 2022;15:800101.

48. Kassabov M, Strogatz SH, Townsend A. A global synchronization theorem for oscillators on a random graph. Chaos: An Interdisciplinary Journal of Nonlinear Science. 2022;32(9).

49. Fix E, Hodges JL. Discriminatory Analysis. Nonparametric Discrimination: Consistency Properties. International Statistical Review / Revue Internationale de Statistique. 1989;57(3):238–47.

50. Yavuz E, Turner J, Nowotny T. GeNN: a code generation framework for accelerated brain simulations. Sci Rep-Uk. 2016;6.

